# Do anthropometric characteristics and physical capacities of highly trained junior badminton players differ according to age and sex?

**DOI:** 10.1101/2025.09.14.676105

**Authors:** Bagus Winata, Joana Brochhagen, Tommy Apriantono, Matthias Wilhelm Hoppe

**Affiliations:** Department of Exercise Science, Institute of Sport Science and Motology, Philipps University of Marburg, Marburg, Germany; Department of Sports Science, School of Pharmacy, Institut Teknologi Bandung, Bandung, Indonesia

## Abstract

The aim of this study was to investigate differences in anthropometric characteristics and physical capacities between (i) under (U) 13, U15, and U17 highly trained junior badminton players and (ii) sexes within each age group. Sixty-two Indonesian highly trained junior badminton players were tested over two sessions for body height, weight, fat mass, and body mass index (BMI), as well as balance, reaction time, hand grip strength, counter movement jump height, linear and non-linear sprint times, and anaerobic sprint and multistage fitness test performances. Traditional (one-way ANOVA or Kruskal-Wallis tests) and alternative statistical approaches (magnitude-based inferences) as well as effect size (ES) calculations were applied for statistical analysis. Regarding age-related differences, in males, U17 players had a statistically significant and most likely higher BMI (*p* = 0.001; ES = very large), as well as statistically significant and most likely superior CMJ height, linear sprint performance, fatigue index, and relative peak power than the U13 players (*p* ≤ 0.003; ES = large to very large). In females, U17 players had a statistically significant and most likely higher BMI and body fat mass (*p* ≤ 0.002; ES = large to very large), as well as statistically significant and very likely inferior non-linear sprint performance and relative peak power than the U13 players (*p* ≤ 0.005; ES = large). For sex-related differences, in U17 and U15 players, males had a statistically significant and most likely lower body fat mass (*p* = 0.001; ES = very large), as well as statistically significant and most likely superior linear sprint performance and relative peak power than females (*p* = 0.001; ES = large to extremely large). Our study shows that anthropometric characteristics and anaerobic capacities differ by age and sex, whereas aerobic capacity is similar among Indonesian highly trained junior badminton players.

## Introduction

Badminton continues to grow with the emergence of numerous competitions across all playing levels [1, 2]. Consequently, many junior players aspire to reach world-class levels, motivating them to participate in long-term developing programs [1]. Primarily, these programs are designed to enhance performance by facilitating training inducted adaptations to meet the physical demands of badminton, typically including repeated short high-intensity actions (e.g., strokes, lunges, jumps, and changes of direction) separated by longer recovery periods at lower intensity (e.g., with a 1:2 work-to-rest ratio) [1–4]. Generally, highly trained junior players start participating in national and international competitions from under (U) 13 to U17. Even at this playing level, match-play characteristics vary not only according to age, but also across the five different playing categories (i.e., men’s and women’s singles as well as men’s, women’s, and mixed doubles), placing specific demands on the players [1, 2]. To meet these demands, a tailored training approach, considering anthropometric characteristics and physical capacities, is essential [1, 2]. Therefore, evaluating both aspects of highly trained junior players according to age and sex using sport-specific testing protocols is a crucial step in optimizing their long-term developments and performances [1, 2], and it requires more research.

In badminton, it is well accepted that anthropometric characteristics, such as body height, weight, and fat mass as well as physical capacities, such as balance, reaction time, strength, speed, and both anaerobic and aerobic endurance, are important prerequisites for meeting the playing demands across all ages and sexes [5–9]. In this context, previous studies have compared anthropometric characteristics and physical capacities between adult elite and sub-elite male players [5, 6]. These studies consistently report that elite players are taller and heavier than their sub-elite counterparts, and demonstrate superior anaerobic and aerobic endurance capacities [5, 6]. However, these differences are reasonably be influenced by training experiences [5, 6]. Considering that existing research has primarily focused on adult male players, a significant research gap remains in highly trained junior players of both sexes [5, 6]. This gap creates ambiguity in designing appropriate training and testing procedures for highly trained junior badminton players. Therefore, evidence-based research in this area is urgently needed to support the long-term development of such players.

To our knowledge, only two studies have compared the anthropometric characteristics and physical capacities of highly trained junior badminton players with those of amateur counterparts [10], while also examining sex-based differences [11]. Although these studies provide valuable insights, they primarily focus on comparisons between different performance levels [10] and investigate a single age group of male and female players [11], leaving age and sex related differences across developmental stages from U13 to U17 players insufficiently explored. Since it is still unclear how these characteristics differ according to age and sex, and given that this period is critical in the development of world-class players [1], it is essential to investigate these differences more comprehensively. Such knowledge may contribute to the development of training programs, specifically tailored to highly trained junior players, and may also assist to design testing protocols as well as support talent identification and player development initiatives [12–14].

Thus, this study aimed to investigate differences in anthropometric characteristics and physical capacities between (i) U13, U15, and U17 highly trained junior badminton players and (ii) sexes within each age group.

## Materials and methods

### Participants

This study included 62 junior male and female badminton players, aged 12-17 years, who participated in national-level junior badminton competitions. The players were from the same badminton club in Indonesia. They had lived in a dormitory since the age of 11, trained for at least 36 hours per week, and competed in at least 14 national tournaments per year. Two U15 male players and four male as well as four female U17 players had prior experience in international-level junior badminton competitions. Based on these attributes, and according to international guidelines, the players were classified as highly trained [2, 15]. All players received an explanation of the study’s purpose, procedures, and potential risks before providing their informed consent. As all players were under the age of 18, written parental consent was obtained via their respective coaches and club committees. All procedures were preapproved by the Ethics Committee of the Bandung Aisyiyah University (1178/kep.01).

According to their ages and sexes, the players were divided into to the following groups: U13 (n = 16; male = 8, female = 8), U15 (n = 20; male = 11, female = 9), and U17 (n = 26; male = 15, female = 11). Players categorized as U13, U15, and U17 had to be born in 2012, 2010, and 2009 or later, respectively, as of August 2024. All dates of birth were obtained and verified by the club committee using players’ birth certificates.

### Experimental study design

A non-interventional cross-sectional study design was used to investigate differences in anthropometric characteristics and physical capacities between the age groups and sexes. The testing procedures were conducted during the second week of the off-season. To ensure maximum performances, players avoided strenuous exercise for 24 hours before testing and stayed well-hydrated. Three physical coaches participated in the testing process, providing motivation and encouragement to the players. The entire session of tests was conducted indoors (temperature: 27–29°C; relative humidity: 78–80%; HTC-2 Clock Temperature, Shenzhen Oway Technology, GD, China) at 07:00 a.m.

The tests were conducted over two sessions. During the first session, anthropometric measures and the following physical capacity tests were performed in the mentioned order: (i) balance test, (ii) reaction test, (iii) hand grip test, (iv) counter movement jump test, (v) linear and non-linear sprint tests, and (vi) anaerobic sprint test. Players were given a 10-minute rest between each test and a 24-hour recovery period before the second testing session. During the latter, an aerobic multistage fitness test was finally conducted.

### Anthropometric measurements

To assess anthropometric characteristics, players were required to wear minimal clothing and be barefoot. Body height was measured using a stadiometer with 0.1 cm readability (Seca 214 Portable Stadiometer, Cardinal Health, Dublin, OH, USA). Furthermore, to assess the percentage of body fat, a 4-point bioelectrical impedance analysis (Omron HBF-375 Body Composition Monitor, Krell Precision, JS, China) with electrodes placed on both hands and feet was utilized, as described previously [16]. The players were asked to stand with their feet on the electrodes, hold the electrode bar, and extend their arms horizontally at a 90° angle from the body. The reliability of the used bioelectric impedance analysis is ICC ≥ 0.90 [17].

### Physical capacity measurements

To assess the dynamic balance ability, a balance test was performed using a digital balance meter (T.K.K. 5,407c Balance-1, Takei Scientific Instrument Co, Niigata, NI, Japan), as applied before [18, 19]. The players were required to stand with one foot on the reaction stand and the other foot on the floor. Upon hearing a signal, the players were required to close both eyes, place their hands on their hips, and lift the foot off the floor until they lost balance. Loss of balance was defined as removing the hands from the hips, lifting the heel or forefoot off the reaction stand, stumbling or falling, or deviating from the testing position. The test was performed twice, with a 1-minute recovery period in between. The maximum time maintained was used for statistical analysis. The reliability of the applied balance test is ICC ≥ 0.90 [20].

To determine the whole-body reaction time, a reaction test using a whole-body reaction measuring equipment (T.K.K. 5,408 Reaction, Takei Scientific Instrument Co, Niigata, NI, Japan) was performed [21]. Reaction was measured using a whole-body reaction measuring equipment (T.K.K. 5,408 Reaction, Takei Scientific Instrument Co, Niigata, NI, Japan). The players were instructed to stand on a contact mat, and upon receiving the visual signal from a flash lamp, to immediately jump off. Reaction time until completely leaving the mat was recorded to assess responsiveness. The test was performed twice, with a 1-minute recovery period in between. The fastest time was used for statistical analysis. The reliability of the used reaction test is ICC ≥ 0.80 [22].

To assess the isometric grip strength, a hand grip test using a digital grip dynamometer (T.K.K. 5,401 Grip-D, Takei Scientific Instrument Co, Niigata, NI, Japan), was performed [23]. The players were instructed to stand upright with their arms hanging by their sides while holding the dynamometer. They were asked to grip it as firmly as possible for 3 seconds without pressing it against their body or bending their elbow. To minimize bias due to intra-individual variabilities, the test was performed four times on both the dominant and non-dominant hand in an alternating manner, with a 1-minute rest period between trials, as recommended by previous studies [24]. The average strength of the four trials for each hand was automatically calculated by the device and used for statistical analysis. The reliability of the hand grip test is ICC ≥ 0.80 [25].

To determine the eccentric-concentric power capacities of the lower extremities, a counter movement jump (CMJ) test using a digital vertical jump meter (T.K.K. 5,414 Jump-DF, Takei Scientific Instrument Co, Niigata, NI, Japan) was performed [26]. The players were instructed to stand on a contact mat with their knees fully extended. Upon hearing a signal, they were required to perform a downward movement followed by a maximal vertical jump with an arm swing [13]. The device calculates vertical jump height based on the flight time. The players performed the CMJ test twice, with a 1-minute recovery period in between. The highest jump was used for statistical analysis. The reliability of the performed CMJ test is ICC ≥ 0.90 [27].

To assess the acceleration, changes of direction, and sprinting capacities, linear and non-linear sprint tests were performed, as described previously [28, 29]. The sprint times were recorded using double-light timing gates (Smart Speed PT Gate System, Fusion Sport, Coopers Plains, QLD, Australia) at the start and finish lines. All players started standing 0.5 meters before the starting line. For the linear sprint test, players were instructed to run as fast as possible for 10 meters [28]. For the non-linear sprint test, the Illinois Test was used, where players were asked to make several changes of direction and finish as quickly as possible, as described in detail elsewhere [29]. Each player was allowed to practice the running pattern before testing. For both sprint tests, the players completed two trials, with a 2-minute rest interval in between. The fastest trial time of each test was used for statistical analysis. The reliability of the used linear and non-linear sprint test is ICC ≥ 0.90 [30, 31].

To determine the anaerobic power and capacity, an anaerobic sprint test was performed, as described in detail before [32]. The sprint times were recorded and controlled with the use of double-light timing gates (Smart Speed PT Gate System, Fusion Sport, Coopers Plains, QLD, Australia) at the start and finish lines over a 35-meter running distance. Briefly, the players were instructed to complete six sprints at maximum effort, with 10 seconds of passive recovery between each sprint [32]. Then, the fatigue index and relative peak power output (PPO) were calculated from the results of the six sprints according to the following formula [32]:

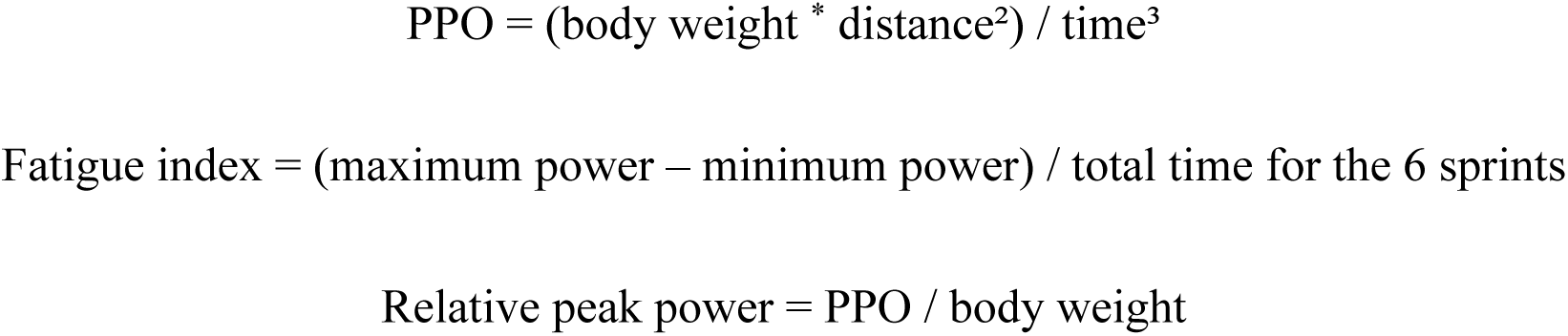

The fatigue index and relative peak power output were used for statistical analysis. The reliability of the applied anaerobic sprint test is ICC ≥ 0.90 [33].

Finally, to assess the aerobic capacities, a multistage fitness test (MFT) was performed, as described in detail before [34]. The total number of shuttles completed by the players during the MFT test was supervised and recorded by the investigators. Briefly, the players were required to perform continuous 20-meter shuttle runs paced by acoustic signals. Each minute, the required running speed was increased. Players were withdrawn from the test, if they failed to maintain the pace for two consecutive shuttles. The MFT results were represented as estimated maximum oxygen uptake (VO_2max_), which was predicted by cross-referencing the final level attained and the number of shuttles completed at the point of exhaustion, as described previously [35]. The VO_2max_ was used for statistical analysis. The reliability of the conducted MFT test is ICC ≥ 0.90 [35].

### Statistical Analysis

Traditional statistical methods were used to examine differences in anthropometric characteristics and physical capacities between ages and sexes. Sample normality and homogeneity were assessed to determine the appropriate statistical tests for each comparison. For comparisons across the three age groups, a one-way ANOVA was applied when assumptions of normality and homogeneity were met; otherwise, the Kruskal-Wallis test was used. To compare differences between both sexes, an independent t-test was employed if the data met the assumptions, and the Mann-Whitney U test was used if they did not. Statistical significance was accepted at *p* < 0.05. To enhance the practical application of our findings, magnitude-based inferences (MBI), as described in detail elsewhere [36, 37], were also calculated. This approach provides a useful statistical alternative by focusing on effect size (ES) and enabling evaluation of meaningfulness, consistent with previous observational studies in applied sport science [12, 13, 38]. The MBI calculations were initially performed by determining the mean values and differences, pooled standard deviations, and ES (Cohen’s d) with 90% confidence intervals (CIs). Subsequently, the positions of the CIs were evaluated in relation to the smallest worthwhile differences (SWDs), defined as the pooled standard deviations multiplied by 0.2, as conducted before [36]. The likelihoods of the differences being truly higher, similar, or lower than the SWD were estimated and interpreted using the following qualitative scale: <1%, most unlikely; 1 to <5%, very unlikely; 5 to <25%, unlikely; 25 to <75%, possibly; 75 to <95%, likely; 95 to <99%, very likely; and ≥99%, most likely. Differences were considered unclear, when there was more than a 5% likelihood of being both higher and lower than the SWD, while only those with a likelihood of ≥75% were deemed practically meaningful [12, 13, 38]. Finally, to clarify the magnitude of the ESs, the following standard thresholds were used: trivial (<0.2), small (0.2 to <0.6), moderate (0.6 to <1.2), large (1.2 to <2.0), very large (2.0 to <4.0), and extremely large (≥4.0) [36].

## Results

Table 1 summarises anthropometric characteristics and physical capacities of the Indonesian highly trained junior badminton players.

**Table 1.** Anthropometric characteristics and physical capacities of Indonesian highly trained junior badminton players.

| Variable | U13 |  | U15 |  | U17 |  |
| --- | --- | --- | --- | --- | --- | --- |
|  | Male (n = 8) | Female (n = 8) | Male (n = 11) | Female (n = 9) | Male (n = 15) | Female (n = 11) |
| Body height (cm) | 158.4 ± 13.5 | 158.2 ± 4.6 | 168.4 ± 5.1 | 156.7 ± 6.5 | 171.1 ± 4.8 | 159.9 ± 3.1 |
| Body weight (kg) | 46.9 ± 13.6 | 49.7 ± 6.2 | 62.5 ± 9.9 | 56.9 ± 4.0 | 66.1 ± 5.8 | 59.9 ± 6.6 |
| BMI (kg/m<sup>2</sup>) | 18.4 ± 2.2 | 19.7 ± 1.8 | 22.0 ± 3.1 | 23.2 ± 1.9 | 22.4 ± 1.7 | 23.3 ± 2.0 |
| Body fat mass (%) | 15.7 ± 4.6 | 22.0 ± 2.7 | 15.3 ± 3.4 | 25.2 ± 2.0 | 15.4 ± 3.7 | 26.6 ± 1.6 |
| Balance (s) | 24.6 ± 23.5 | 46.5 ± 17.6 | 39.4 ± 30.5 | 52.8 ± 28.1 | 39.0 ± 34.1 | 31.3 ± 25.6 |
| Reaction (s) | 0.3 ± 0.0 | 0.7 ± 1.0 | 0.3 ± 0.0 | 0.3 ± 0.0 | 0.3 ± 0.0 | 0.3 ± 0.0 |
| Dominant hand grip (kgf) | 28.0 ± 9.9 | 26.5 ± 4.2 | 41.7 ± 6.1 | 32.2 ± 2.9 | 42.6 ± 4.7 | 29.4 ± 3.9 |
| Non-dominant hand grip (kgf) | 24.6 ± 7.5 | 21.8 ± 3.9 | 32.3 ± 7.4 | 25.6 ± 2.1 | 34.9 ± 3.4 | 22.6 ± 3.1 |
| CMJ (cm) | 50.8 ± 4.5 | 46.4 ± 3.9 | 57.9 ± 6.1 | 46.0 ± 3.1 | 61.1 ± 3.5 | 45.2 ± 4.1 |
| Linear sprint (s) | 2.0 ± 0.1 | 2.1 ± 0.1 | 1.8 ± 0.1 | 2.1 ± 0.1 | 1.8 ± 0.1 | 2.1 ± 0.1 |
| Non-linear sprint (s) | 17.7 ± 1.2 | 18.2 ± 0.5 | 17.5 ± 0.8 | 18.4 ± 0.9 | 17.9 ± 2.7 | 19.5 ± 1.1 |
| Fatigue index (W/s) | 2.7 ± 1.3 | 3.4 ± 1.4 | 5.4 ± 2.0 | 3.2 ± 1.2 | 6.9 ± 2.1 | 2.6 ± 1.0 |
| Relative peak power (W/kg) | 6.4 ± 1.6 | 6.8 ± 1.3 | 8.1 ± 1.2 | 6.0 ± 0.8 | 9.7 ± 1.2 | 5.5 ± 0.7 |
| VO<sub>2max</sub> (ml/kg/min) | 51.6 ± 3.0 | 50.0 ± 4.8 | 52.2 ± 8.0 | 50.6 ± 7.6 | 50.5 ± 5.6 | 51.5 ± 5.7 |

Note: U13, U15, and U17 = under 13, 15 and 17 years old players; n = sample size; BMI = body mass index; CMJ = counter movement jump.

### Differences between the age groups

Table 2 presents the results of traditional statistical analyses, examining differences in anthropometric characteristics and physical capacities between ages among Indonesian highly trained junior badminton players. Figures 1 and 2 illustrate the results of the alternative statistical approach used to explore these differences in male and female players, respectively. In males, the U17 players had a statistically significant and most likely higher BMI than the U13 players (*p* = 0.001; ES = very large). The U15 players had a statistically significant and likely higher BMI than the U13 players (*p* = 0.016; ES = large). The U17 players had a statistically significant and very likely higher relative peak power than the U15 players (*p* = 0.010; ES = large). The U17 players had a statistically significant and possibly superior dominant hand grip strength, likely superior non-dominant hand grip strength, and most likely superior CMJ height, linear sprint performance, fatigue index, and relative peak power than the U13 players (*p* ≤ 0.003; ES = large to very large). The U15 players had a statistically significant and likely higher relative peak power as well as very likely superior linear sprint performance and fatigue index than the U13 players (*p* ≤ 0.035; ES = moderate to large).

**Figure 1.**
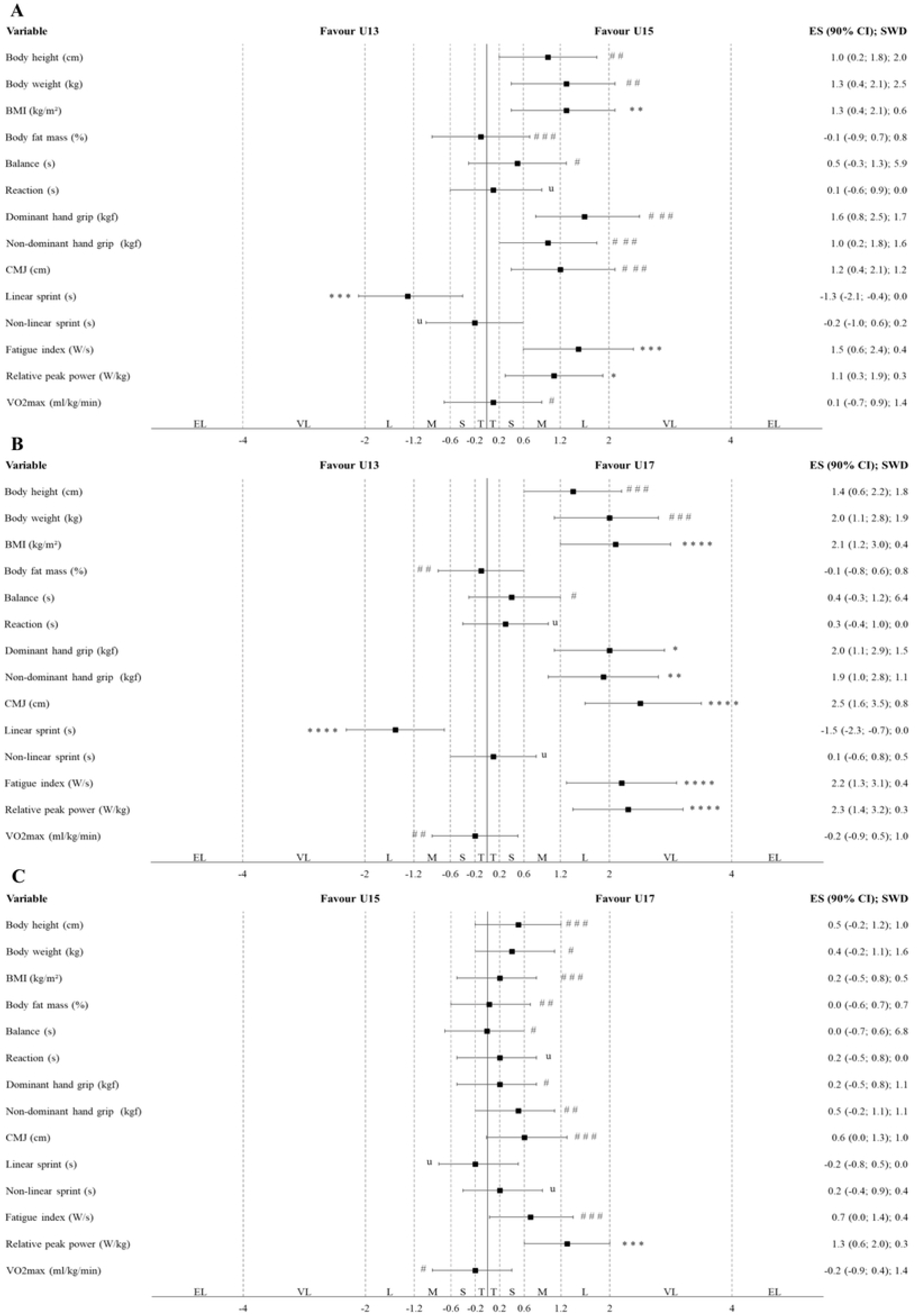
Differences in anthropometric characteristics and physical capacities between the different age groups in male Indonesian highly trained junior badminton players (A: U13 vs. U15; B: U13 vs. U17; C: U15 vs. U17) according to the alternative statistical approaches (magnitude-based inferences). Note: The CIs of ESs were evaluated in relation to the SWDs. U13, U15, and U17 = under 13, 15 and 17 years old players; BMI = body mass index; CMJ = counter movement jump; T = trivial; S = small; M = moderate; L = large; VL = very large; EL = extremely large. The symbols #, ##, and ### indicate the probability that the effect is higher and lower than the SWD: <1% (most unlikely), <5% (very unlikely), and <74% (unlikely), respectively. Meanwhile, *, **, ***, and **** indicate the probability that the effect is higher or lower than the SWD: >75% (possibly), >90% (likely), >95.5% (very likely), and 100% (most likely), respectively. The letter “u” denotes an unclear effect, with a probability of both higher and lower than 5% in either direction relative to the SWD.

**Figure 2.**
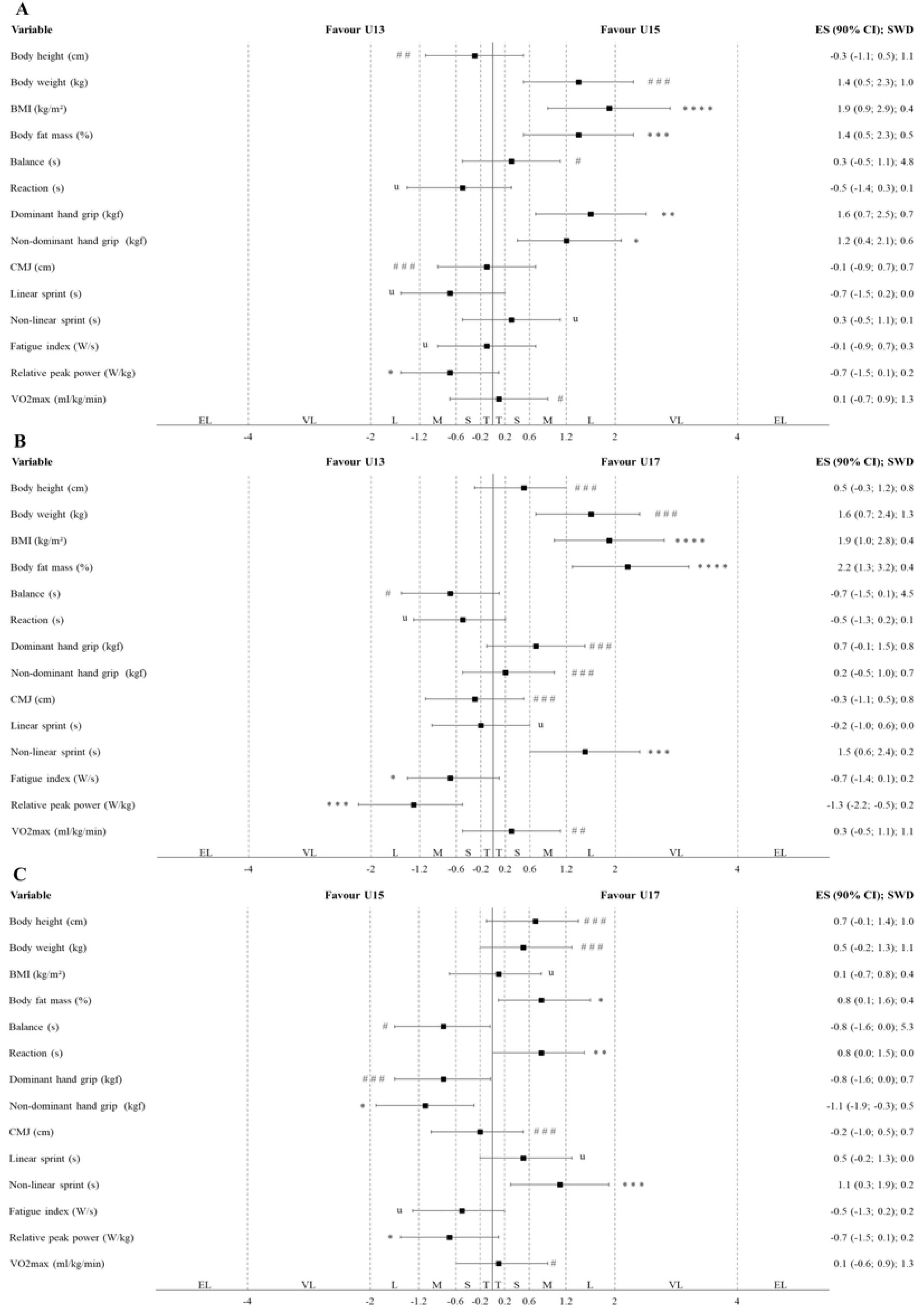
Differences in anthropometric characteristics and physical capacities between the different age groups in female Indonesian highly trained junior badminton players (A: U13 vs. U15; B: U13 vs. U17; C: U15 vs. U17) according to the alternative statistical approaches (magnitude-based inferences). Note: The CIs of ESs were evaluated in relation to the SWDs. U13, U15, and U17 = under 13, 15 and 17 years old players; BMI = body mass index; CMJ = counter movement jump; T = trivial; S = small; M = moderate; L = large; VL = very large; EL = extremely large. The symbols #, ##, and ### indicate the probability that the effect is higher and lower than the SWD: <1% (most unlikely), <5% (very unlikely), and <74% (unlikely), respectively. Meanwhile, *, **, ***, and **** indicate the probability that the effect is higher or lower than the SWD: >75% (possibly), >90% (likely), >95.5% (very likely), and 100% (most likely), respectively. The letter “u” denotes an unclear effect, with a probability of both higher and lower than 5% in either direction relative to the SWD.

**Table 2.** Differences in anthropometric characteristics and physical capacities between the different age groups and both sexes of Indonesian highly trained junior badminton players according to the traditional statistical approaches (one-way ANOVA or Kruskal-Wallis tests).

| Variable | Male players |  |  |  | Female players |  |  |  |
| --- | --- | --- | --- | --- | --- | --- | --- | --- |
|  | One-way ANOVA or Kruskal-Wallis tests (*p*-value) | Bonferroni or Pairwise tests (*p*-value) |  |  | One-way ANOVA or Kruskal-Wallis tests (*p*-value) | Bonferroni or Pairwise tests (*p*-value) |  |  |
|  |  | U13 vs. U15 | U13 vs. U17 | U15 vs. U17 |  | U13 vs. U15 | U13 vs. U17 | U15 vs. U17 |
| Body height (cm) | 0.026 * | 0.097 | 0.007 * | 0.300 | 0.409 | 1.000 | 1.000 | 0.457 |
| Body weight (kg) | 0.005 * | 0.063 | 0.001 * | 0.155 | 0.003 * | 0.017 * | 0.001 * | 0.374 |
| BMI (kg/m<sup>2</sup>) | 0.003 * | 0.016 * | 0.001 * | 0.364 | 0.002 * | 0.003 * | 0.002 * | 0.976 |
| Body fat mass (%) | 0.973 | 1.000 | 1.000 | 1.000 | 0.001 * | 0.011 * | 0.001 * | 0.384 |
| Balance (s) | 0.559 | 0.934 | 0.885 | 1.000 | 0.121 | 1.000 | 0.580 | 0.187 |
| Reaction (s) | 0.789 | 1.000 | 1.000 | 1.000 | 0.125 | 0.499 | 0.519 | 1.000 |
| Dominant hand grip (kgf) | 0.004 * | 0.012 * | 0.001 * | 0.541 | 0.019 * | 0.006 * | 0.323 | 0.052 |
| Non-dominant hand grip (kgf) | 0.002 * | 0.028 * | 0.001 * | 0.821 | 0.038 * | 0.049 * | 1.000 | 0.117 |
| CMJ (cm) | 0.001 * | 0.008 * | 0.001 * | 0.283 | 0.777 | 1.000 | 1.000 | 1.000 |
| Linear sprint (s) | 0.002 * | 0.012 * | 0.003 * | 1.000 | 0.338 | 0.476 | 1.000 | 0.878 |
| Non-linear sprint (s) | 0.790 | 1.000 | 1.000 | 1.000 | 0.008 * | 0.672 | 0.005 * | 0.015 * |
| Fatigue index (W/s) | 0.001 * | 0.012 * | 0.001 * | 0.170 | 0.338 | 1.000 | 0.517 | 0.849 |
| Relative peak power (W/kg) | 0.001 * | 0.035 * | 0.001 * | 0.010 * | 0.014 * | 0.228 | 0.004 * | 0.094 |
| VO<sub>2max</sub> (ml/kg/min) | 0.581 | 1.000 | 1.000 | 1.000 | 0.856 | 1.000 | 1.000 | 1.000 |

Note: U13, U15, and U17 = under 13, 15 and 17 years old players; BMI = body mass index; CMJ = counter movement jump; \* = significance level indicated at $p < 0.05$ .

Meanwhile, in females, the U17 players had a statistically significant and most likely higher BMI and body fat mass than the U13 players (*p* ≤ 0.002; ES = large to very large). The U15 players had a statistically significant and very likely higher body fat mass as well as most likely higher BMI than the U13 players (*p* ≤ 0.011; ES = large). The U17 players had a statistically significant and very likely inferior non-linear sprint performance than the U15 players (*p* = 0.015; ES = moderate). The U17 players had a statistically significant and very likely inferior non-linear sprint performance and relative peak power than the U13 players (*p* = ≤ 0.005; ES = large). The U15 players had a statistically significant and possibly superior non-dominant hand grip strength and likely superior dominant hand grip strength than the U13 players (*p* ≤ 0.049; ES = large).

### Differences between the sexes

Table 3 and Figure 3 present the results of traditional and alternative statistical analyses, respectively, used to examine differences in anthropometric characteristics and physical capacities between sexes within each age group of Indonesian highly trained junior badminton players. In U17 players, males had a statistically significant and most likely higher body height, and lower body fat mass than females (*p* = 0.001; ES = very large). In U15 players, males had a statistically significant and likely higher body height as well as most likely lower body fat mass than females (*p* = 0.001; ES = very large). In U13 players, males had a statistically significant and likely lower body fat mass than females (*p* = 0.005; ES = large). In U17 players, males had a statistically significant and possibly superior non-linear sprint performance as well as most likely superior reaction time, dominant hand grip strength, non-dominant hand grip strength, CMJ height, linear sprint performance, fatigue index, and relative peak power than females (*p* ≤ 0.006; ES = moderate to extremely large). In U15 players, males had a statistically significant and very likely superior dominant hand grip strength, CMJ height, non-linear sprint performance, and fatigue index as well as most likely superior linear sprint performance and relative peak power than females (*p* ≤ 0.021; ES = moderate to very large). In U13 players, males had a statistically significant and most likely superior linear sprint performance than females (*p* = 0.009; ES = large).

**Figure 3.**
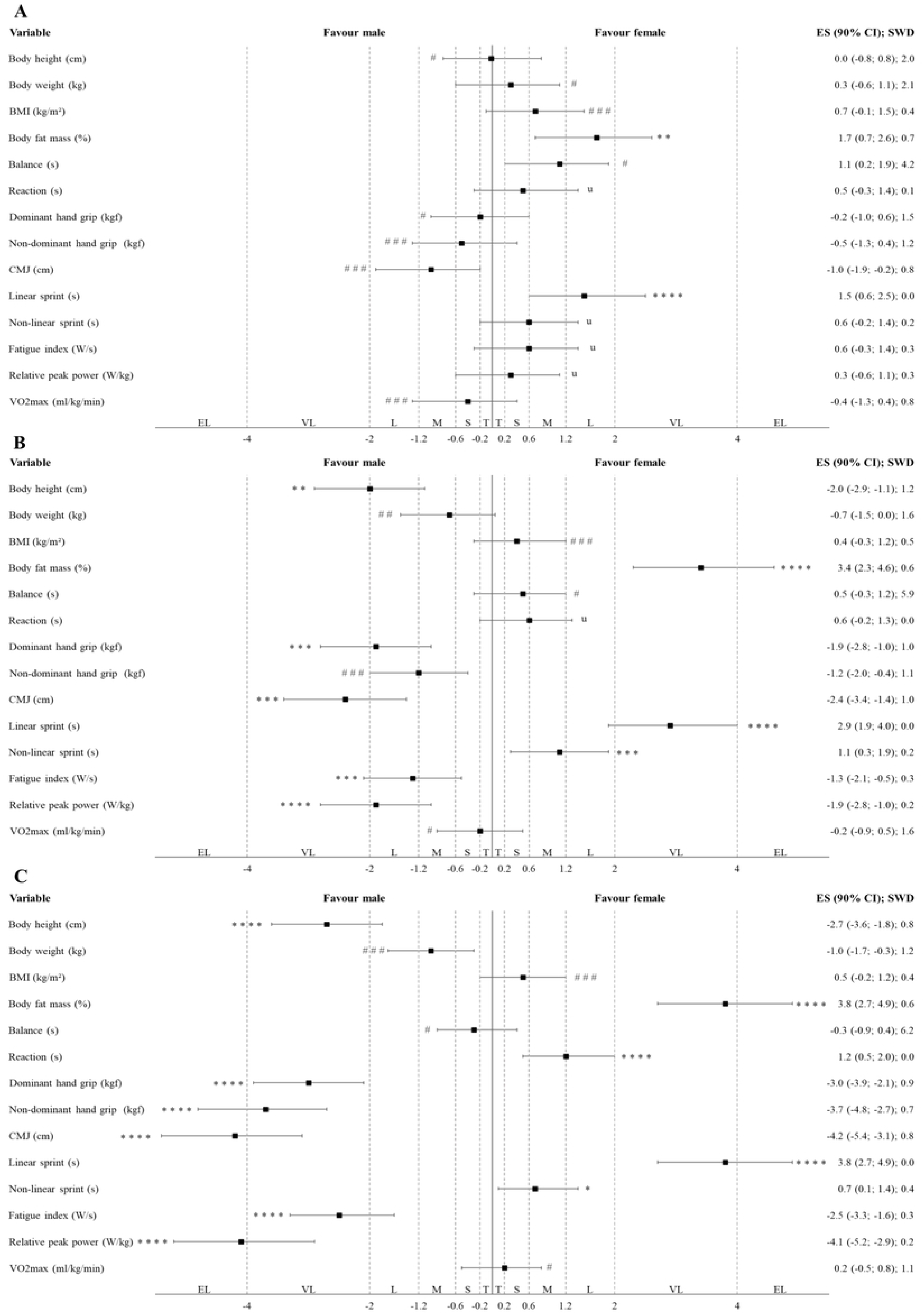
Differences in anthropometric characteristics and physical capacities between sexes within each age group of Indonesian highly trained junior badminton players (A: U13; B: U15; C: U17) according to alternative statistical approaches (magnitude-based inferences). Note: The CIs of ESs were evaluated in relation to the SWDs. U13, U15, and U17 = under 13, 15 and 17 years old players; BMI = body mass index; CMJ = counter movement jump; T = trivial; S = small; M = moderate; L = large; VL = very large; EL = extremely large. The symbols #, ##, and ### indicate the probability that the effect is higher and lower than the SWD: <1% (most unlikely), <5% (very unlikely), and <74% (unlikely), respectively. Meanwhile, *, **, ***, and **** indicate the probability that the effect is higher or lower than the SWD: >75% (possibly), >90% (likely), >95.5% (very likely), and 100% (most likely), respectively. The letter “u” denotes an unclear effect, with a probability of both higher and lower than 5% in either direction relative to the SWD.

**Table 3.** Differences in anthropometric characteristics and physical capacities between both sexes of Indonesian highly trained junior badminton players according to the traditional statistical approaches (independent t-test or Mann–Whitney U test).

| Variable | U13 | U15 | U17 |
| --- | --- | --- | --- |
|  | Independent t-test<br>or Mann-Whitney U test<br>(*p*-value) | Independent t-test<br>or Mann-Whitney U test<br>(*p*-value) | Independent t-test<br>or Mann-Whitney U test<br>(*p*-value) |
| Body height (cm) | Males (0.834) | Males (0.001) * | Males (0.001) * |
| Body weight (kg) | Females (0.601) | Males (0.138) | Males (0.021) * |
| BMI (kg/m<sup>2</sup>) | Females (0.183) | Females (0.080) | Females (0.436) |
| Body fat mass (%) | Females (0.005) * | Females (0.001) * | Females (0.001) * |
| Balance (s) | Females (0.045) * | Females (0.325) | Males (0.421) |
| Reaction (s) | Females (0.006) * | Females (0.204) | Females (0.004) * |
| Dominant hand grip (kgf) | Males (0.834) | Males (0.001) * | Males (0.001) * |
| Non-dominant hand grip (kgf) | Males (0.375) | Males (0.012) * | Males (0.001) * |
| CMJ (cm) | Males (0.058) | Males (0.001) * | Males (0.001) * |
| Linear sprint (s) | Females (0.009) * | Females (0.001) * | Females (0.001) * |
| Non-linear sprint (s) | Females (0.115) | Females (0.021) * | Females (0.006) * |
| Fatigue index (W/s) | Females (0.269) | Males (0.010) * | Males (0.001) * |
| Relative peak power (W/kg) | Females (0.613) | Males (0.001) * | Males (0.001) * |
| VO<sub>2max</sub> (ml/kg/min) | Males (0.413) | Males (0.658) | Females (0.835) |

Note: “Males” or “Females” indicates that the variable had a higher value in the specified sex. U13, U15, and U17 = under 13, 15 and 17 years old players; BMI = body mass index; CMJ = counter movement jump; \* = significance level indicated at $p < 0.05$ .

## Discussion

This study is the first to investigate differences in anthropometric characteristics and physical capacities between (i) U13, U15, and U17 Indonesian highly trained junior badminton player and (ii) sexes within each age group. Our main findings were: (i) the age differences showed that, in males, U17 and U15 players had higher BMI and showed superior performance in tests related to anaerobic capacities compared to U13 players, whereas in females, U17 and U15 players had higher BMI and body fat mass than U13 players, but U17 players showed lower anaerobic capacities compared to both U15 and U13 players; and (ii) the largest sex differences were observed in U17 and U15 players, with males showing higher body height and superior performance in tests related to anaerobic capacities and females exhibiting higher body fat mass.

This study provides new knowledge into the anthropometric characteristics and physical capacities of junior badminton players, addressing limitations in previous studies that mainly compared these variables between highly trained juniors and amateur players [10] as well as between sexes [11] only. The most notable differences by age and sex were observed in anthropometric characteristics and anaerobic capacities (Table 1). Regarding age-related differences, our findings suggest that growth and maturation processes are most likely the primary factors underlying differences in anthropometric characteristics and anaerobic capacities in badminton, similar with patterns reported in other sports such as soccer [12] and handball [13]. Furthermore, our findings on sex-related differences may help explain the variations in match-play characteristics reported in a previous systematic review [2]. Specifically, that review stated that male players tend to perform more explosive and high-intensity movements than female players [2]. This aligns with our results, which show that males have greater body height, lower body fat, and superior anaerobic capacities compared with females, which may enable them to execute more explosive and high-intensity movements. In contrast, female players, with shorter body height, higher body fat, and lower anaerobic capacities, may be more limited in producing such actions and thus tend to exhibit more defensive match-play characteristics.

Our first main finding was that, in males, highly trained Indonesian junior badminton players in the U17 and U15 players had higher BMI (ES = large to very large) and superior performance in tests related to anaerobic capacities (ES = moderate to very large) than U13 players, whereas in females, U17 and U15 had higher BMI and body fat mass (ES = large to very large) than U13 players, but U17 players showed inferior performance in tests related to anaerobic capacities (ES = moderate to large) compared to both U15 and U13 players (Table 2, Figure 1–2). Our results are supported by two narrative reviews [1, 7], which found that among players aged U13 to U17, older junior badminton players, particularly males, had higher BMI [1, 7] and superior CMJ [7] and higher relative peak power [7], while older females showed higher BMI [1,7] and body fat mass [1,7] than their younger counterparts. A possible explanation for our findings is that growth and maturation processes start around this age [12, 39–42]. For example, U17 male players typically have more mature testosterone development compared to U15 and U13 players [42]. This hormonal difference contributes to increased muscle mass, which may allow for them to produce superior performance in tests related to anaerobic capacities [12, 42, 43]. Conversely, in females, U17 players tend to have more mature oestrogen development, which is associated with increased body fat mass compared to U15 and U13 females, and relatively smaller muscle size compared to males [42]. As a result, U17 female players may show inferior performance in tests related to anaerobic capacities [42, 43]. Meanwhile, our findings also revealed that VO_2max_ levels were relatively similar among U13 to U17 players in both sexes, with values ranging from 50.5–52.2 ml/kg/min in males and 50.0–51.5 ml/kg/min in females (Table 1). Unfortunately, no observational study specifically comparing estimated or measured VO_2max_ levels according to different age groups in badminton is currently available. However, in others sports like soccer, studies indicate that VO_2max_ levels of junior players have remained unchanged over the past decade [12, 44]. Future studies in badminton are needed to address this research gap. Overall, these findings suggest that age differences in both sexes are primarily associated with variations in anthropometric characteristics and anaerobic capacity among Indonesian highly trained junior badminton players.

Our second main finding was that the largest sex differences among Indonesian highly trained junior badminton players were observed in the U17 and U15 players, in contrast to the U13 players. Specifically, U17 and U15 male players showed significantly higher body height (ES = very large), lower body fat mass (ES = very large), and superior performance in tests again related to anaerobic capacities (ES = moderate to extremely large) compared to female players (Table 3, Figure 3). These results are consistent with previous observational studies, examining sex differences in anthropometric characteristics and physical capacities, particularly in U17 players [11]. Although studies focusing specifically on junior badminton players are limited, similar trends have been reported in studies involving adult players [45–47]. Differences in peak height velocity (PHV) between males and females may help explain these findings [12, 39, 40]. A previous study indicates that PHV typically occurs later in males (13 to 15 years) than in females (11 to 13 years) [39]. During PHV, males experience a rapid increase in body height and muscle mass, accompanied by a relative decrease in body fat mass [12, 39, 40]. In contrast, while females also experience growth in stature, this phase is often associated with an increase in body fat mass [12, 39, 40]. These anthropometric characteristic differences substantially influence anaerobic capacities in both sexes, as supported by prior studies [12, 39, 40, 42].

Additionally, our second finding also indicated that female Indonesian highly trained junior badminton players had VO_2max_ levels that were similar to their male counterparts across all age groups (U13 = 50.0 vs. 51.6; U15 = 50.6 vs. 52.2; U17 = 51.5 vs. 50.5 ml/kg/min, respectively), as shown in Table 1. This finding contrasts with a previous review reporting lower VO_2max_ levels in female junior players compared to their male counterparts (48.1 vs. 57.2 ml/kg/min) [1]. One possible explanation for this finding is the influence of factors such as competitive participation [12, 48] and specific training adaptations targeting the cardiovascular and neuromuscular systems [12, 49]. In fact, these Indonesian highly trained junior players have been engaged in systematic training programs since the age of 11 at a dedicated badminton academy. These programs are designed to be intensive and specifically tailored to the physical and tactical demands of their match play category. For example, to meet the demands of competitive play, which in female players often involves more defensive play and smoother shots, training programs tend to emphasize the development and maintenance of aerobic capacity [2]. As a result, long-term adaptations tend to support relatively higher VO_2max_ levels [50, 51]. In contrast, the match play of males, characterized by more explosive and high-intensity movements, leads to training that prioritizes anaerobic capacity [2]. This results in more pronounced long-term adaptations in strength and power, but less emphasis on improvements in VO_2max_ levels [50, 51]. Overall, these findings suggest that sex differences within U17 and U15 age groups are associated with variations in anthropometric characteristics and anaerobic capacity among Indonesian highly trained junior badminton players. However, the factors assumed to influence these differences (PHV, competitive participation, and specific training adaptations) should be interpreted with caution, as they were not directly measured in this study.

Overall, this study provides first evidence of differences in the anthropometric characteristics and physical capacities of Indonesian highly trained junior badminton players, which are relatively similar to findings in other racquet sports, such as tennis [52]. From a practical perspective, training programs for these players should be carefully tailored to account for variations in anthropometric characteristics and physical capacities according to age and sex. For instance, within the U13 to U17 age range in both sexes, U17 male players should focus on optimizing anaerobic capacity more than U15 and U13 players, while U15 males should focus more than U13 males. Similarly, U17 female players may benefit most from developing aerobic capacity, followed by U15 and then U13 players. The implementation of intermittent training drills, designed with specific metabolic work durations (e.g., less than 30 seconds to stimulate anaerobic systems and more than 2 minutes to engage aerobic systems), can serve as a relevant example of such training programs. Concerning testing programs, our findings indicate that anaerobic power-oriented testing is more appropriate for male players, whereas aerobic endurance testing may be more beneficial for evaluating female players. Lastly, our results may serve as a foundational framework for talent identification and player development initiatives, aiming to guide players toward achieving world-class performance levels.

We acknowledge several limitations of this study. First, the cross-sectional design does not allow for strong conclusions regarding the influence of growth and maturation processes, competitive participation, or specific training adaptations. Therefore, a longitudinal study is required to more accurately address this research gap. Second, VO_2max_ was not directly measured, but rather estimated using the well-established MFT. Therefore, further studies are needed to address these issues.

## Conclusion

Our study shows that anthropometric characteristics and anaerobic capacities differ by age and sex, whereas aerobic capacity is similar among Indonesian highly trained junior badminton players. Thus, the development of training and testing programs, as well as a framework for talent identification for highly trained junior badminton players, must carefully account for these differences to support their progression toward world-class levels.

## Acknowledgement

The authors would like to thank Muhamad Fahmi Hasan, Agung Dwi Juniarsyah, and members of the Sports Science Department at Institut Teknologi Bandung for their assistance during data collection. The authors also gratefully acknowledge the support of the Badminton World Federation (BWF) for this study. Open access funding provided by the Open Access Publishing Fund of Philipps-Universität Marburg.

## Supporting information

**S1 Figure 1. Differences in anthropometric characteristics and physical capacities between the different age groups in male Indonesian highly trained junior badminton players (A: U13 vs. U15; B: U13 vs. U17; C: U15 vs. U17) according to the alternative statistical approaches (magnitude-based inferences).**

Note: The CIs of ESs were evaluated in relation to the SWDs.

U13, U15, and U17 = under 13, 15 and 17 years old players; BMI = body mass index; CMJ = counter movement jump; T = trivial; S = small; M = moderate; L = large; VL = very large; EL = extremely large. The symbols #, ##, and ### indicate the probability that the effect is higher and lower than the SWD: <1% (most unlikely), <5% (very unlikely), and <74% (unlikely), respectively. Meanwhile, *, **, ***, and **** indicate the probability that the effect is higher or lower than the SWD: >75% (possibly), >90% (likely), >95.5% (very likely), and 100% (most likely), respectively. The letter “u” denotes an unclear effect, with a probability of both higher and lower than 5% in either direction relative to the SWD.

**S2 Figure 2. Differences in anthropometric characteristics and physical capacities between the different age groups in female Indonesian highly trained junior badminton players (A: U13 vs. U15; B: U13 vs. U17; C: U15 vs. U17) according to the alternative statistical approaches (magnitude-based inferences)**.

Note: The CIs of ESs were evaluated in relation to the SWDs.

U13, U15, and U17 = under 13, 15 and 17 years old players; BMI = body mass index; CMJ = counter movement jump; T = trivial; S = small; M = moderate; L = large; VL = very large; EL = extremely large. The symbols #, ##, and ### indicate the probability that the effect is higher and lower than the SWD: <1% (most unlikely), <5% (very unlikely), and <74% (unlikely), respectively. Meanwhile, *, **, ***, and **** indicate the probability that the effect is higher or lower than the SWD: >75% (possibly), >90% (likely), >95.5% (very likely), and 100% (most likely), respectively. The letter “u” denotes an unclear effect, with a probability of both higher and lower than 5% in either direction relative to the SWD.

**S3 Figure 3. Differences in anthropometric characteristics and physical capacities between sexes within each age group of Indonesian highly trained junior badminton players (A: U13; B: U15; C: U17) according to alternative statistical approaches (magnitude-based inferences).**

Note: The CIs of ESs were evaluated in relation to the SWDs.

U13, U15, and U17 = under 13, 15 and 17 years old players; BMI = body mass index; CMJ = counter movement jump; T = trivial; S = small; M = moderate; L = large; VL = very large; EL = extremely large. The symbols #, ##, and ### indicate the probability that the effect is higher and lower than the SWD: <1% (most unlikely), <5% (very unlikely), and <74% (unlikely), respectively. Meanwhile, *, **, ***, and **** indicate the probability that the effect is higher or lower than the SWD: >75% (possibly), >90% (likely), >95.5% (very likely), and 100% (most likely), respectively. The letter “u” denotes an unclear effect, with a probability of both higher and lower than 5% in either direction relative to the SWD.

**S1 Table 1. Anthropometric characteristics and physical capacities of Indonesian highly trained junior badminton players.**

Note: U13, U15, and U17 = under 13, 15 and 17 years old players; n = sample size; BMI = body mass index; CMJ = counter movement jump.

**S2 Table 2. Differences in anthropometric characteristics and physical capacities between the different age groups and both sexes of Indonesian highly trained junior badminton players according to the traditional statistical approaches (one-way ANOVA or Kruskal-Wallis tests).**

Note: U13, U15, and U17 = under 13, 15 and 17 years old players; BMI = body mass index; CMJ = counter movement jump; * = significance level indicated at p < 0.05.

**S3 Table 3. Differences in anthropometric characteristics and physical capacities between both sexes of Indonesian highly trained junior badminton players according to the traditional statistical approaches (independent t-test or Mann–Whitney U test).**

Note: “Males” or “Females” indicates that the variable had a higher value in the specified sex. U13, U15, and U17 = under 13, 15 and 17 years old players; BMI = body mass index; CMJ = counter movement jump; * = significance level indicated at p < 0.05.

